# Genes on the Move: Exploring the Resistome of Viable Isolates from a First Nation Community in Manitoba

**DOI:** 10.1101/2025.07.11.664311

**Authors:** Rudra Patel, Jocelyn Zambrano-Alvarado, Nick Rudin, Karl Zhang, Adetola Adeniji, M. Moniruzzaman, Miguel Uyaguari-Diaz

## Abstract

Despite wastewater treatment plants (WWTPs) and oxidation lagoons (OL) being designed for pollutant and pathogen removal, these systems can also serve as environments conducive to the exchange of mobile genetic elements (MGEs) and antibiotic resistance genes (ARGs) among microorganisms. This context facilitates the dissemination of antimicrobial resistance (AMR) among both pathogenic and non-pathogenic environmental bacteria, as well as phage populations, within these facilities. To enhance understanding of AMR dynamics in wastewater, our study examined the resistome of bacterial isolates recovered from a OL in a First Nation (FN) community in Manitoba, Canada. Thirty-five samples were collected from all OL stages, including raw sewage, lagoons, submerged attached growth reactor (SAGR), UV-treated effluent, and an upstream control site, from September 2022 to April 2023. Whole genome sequencing of 58 viable isolates was performed using Oxford Nanopore MinION, with genomes reconstructed via *de novo* assembly. The isolates corresponded to the following genera: *Aeromonas* 50.87% *, Serratia* 15.78%, *Pantoea* and *Escherichia* 7.01% each*, Lelliottia* 5.26%*, Rahnella, Enterobacter* and *Buttiauxella* 3.50% each, and finally *Acinetobacter, Yersinia* and *Citrobacter* 1.75%. Furthermore, a total of 32,559 elements were classified as ARGs. MGEs carrying ARGs were detected in 72.72*%* of the isolates. ARGs conferring resistance to macrolides, tetracyclines, and fluoroquinolones demonstrated a notable presence within MGEs (plasmids and bacteriophages) identified throughout the facility’s stages. This research represents a non-invasive approach, enabling an in-depth exploration of AMR dynamics by discerning the resistome profiles of bacteria residing within a rural OL facility from a FN community.

**Importance:** OLs commonly found in rural and Indigenous communities in Canada, are crucial for mitigating environmental hazards and safeguarding public health. Paradoxically, these systems can also serve as hotspots for the dissemination of ARGs and MGEs, thereby exacerbating AMR. The persistence of ARGs within both pathogenic and non-pathogenic bacteria, often carried on MGEs including phages, in treated effluents raises significant concerns about the environmental impact and public health. This study offers fundamental insights into the unique dynamics of AMR within OLs serving a First Nation community in Manitoba. Understanding these specific dynamics is crucial for developing robust monitoring frameworks and implementing targeted, sustainable strategies to mitigate the public health and environmental risks posed by AMR in these often-underserved remote settings.

## Introduction

Wastewater treatment plants (WWTPs) and oxidation lagoons (OLs) are intricate infrastructures engineered to receive, concentrate, and treat organic waste (1). Their primary objective is the removal of pollutants and pathogens present in wastewater, a crucial step in mitigating potential environmental hazards and safeguarding public health. OLs function as small, self-sufficient ponds, leveraging microorganisms, light, and the presence of algae for the removal of organics and other contaminants (2). OLs are recognized for their cost-effectiveness (3) and constitute a prevalent method for wastewater treatment in rural and First Nation (FN) communities across Canada (4). In Manitoba alone, there are approximately 350 OLs (4). While WWTPs and OLs play a pivotal role in maintaining public health by treating wastewater effectively, it has been documented that they also serve as reservoirs for antibiotic resistance bacteria (ARB) (1, 3). As a consequence of the widespread use of antimicrobial agents, disinfectants, and antibiotics, especially after the COVID-19 pandemic, antibiotic-resistant bacteria (ARB) and pathogens are consistently discharged into the environment (5). These environmental reservoirs of ARB, antibiotics, and heavy metals resulting from anthropogenic activities subsequently impact the composition of WWTPs and OLs influents. OLs rely mainly on primary physical treatment, such as aeration, sunlight, and gravity, for the clarification, sedimentation, and subsequent breakdown of organic waste present in wastewater (2). The complex microbial communities within OLs, coupled with the selective pressures exerted during treatment processes, create an environment conducive to the survival and propagation of ARB and antibiotic resistance genes (ARGs) (6, 7). Additionally, having steps in the process that span several weeks facilitates bacterial conjugation (9). The persistence of these antibiotics and genetic elements underscores the complexity of wastewater treatment and raises concerns about the potential environmental impact of treated effluents.

Mobile genetic elements (MGEs), including plasmids, transposons, and integrons, are extensively characterized genetic components crucial for the transfer of ARGs through mechanisms such as conjugation and transformation (11). Bacteriophages or phages, which are viruses that infect bacteria, have recently emerged as another noteworthy source of ARGs capable of facilitating resistance transfer through a process known as transduction (13). The mobilome (collection of all MGEs)(6) and the recognition of multiple pathways for the dissemination of antibiotic resistance highlight the dynamic and multifaceted nature of the resistome (the collection of all ARGs found in pathogenic and non-pathogenic bacteria) (7, 8) within microbial communities. This resistome represents a threat toward the diminishing efficacy of antibiotics as a therapeutic option for treating bacterial infections (12). To avert this impending crisis and preserve the efficacy of antibiotics, it becomes imperative to address the root cause by mitigating the spread of antimicrobial resistance (AMR). OLs serve as open-system models, which may facilitate the examination of genetic profiles within isolated microbial communities obtained from water samples. Furthermore, studies characterizing the resistome in OLs in FN communities in Canada and North America remain scarce. The specific OLs investigated in this study were associated with the wastewater treatment facility of a FN community with a population <5,000 inhabitants, and situated two hours radius of Winnipeg, MB. Notably, the OLs in this FN are integrated with a Submerged Attached Growth Reactor (also known as SAGR), a design developed based on the concept of an aerated gravel bed (9). This integrated system enhances treatment efficiency and underscores the innovative approaches implemented in wastewater management within such community contexts. The SAGR incorporates a distinctive feature in which the gravel bed serves as a mechanism to trap and retain microorganisms. This design prevents the inadvertent washout of microorganisms into the effluent, ensuring their sustained presence within the system (3). Between September 2022 and April 2023, a monthly sampling regimen was implemented, encompassing the spectrum from the most contaminated source, raw sewage (RS), to the least contaminated source, treated effluent (EF) after UV irradiation. This systematic sampling approach aimed at evaluating the natural flow of wastewater within the system. In the OLs, RS progresses from lagoons to SAGR beds and is subsequently exposed to UV treatment. In this context, samples collected from lagoons represent the pre-SAGR stage, while those from the SAGR denote the post-SAGR filtration stage. Following treatment of SAGR, wastewater undergoes UV before being discharged as effluent into the aquatic environment.

To provide a comprehensive understanding of microbial dynamics, a case-control study was conducted by collecting water samples from a location upstream (UP) of the FN community and wastewater effluent site. This strategic sampling aimed to capture baseline microbial profiles unaffected by the treatment processes, enabling a comparative analysis across different stages of wastewater treatment. Through a detailed examination of the genetic profiles, resistome, and mobilome within OLs, this study provides fundamental insights into the critical roles of wastewater treatment in the dissemination of AMR of a FN community. A deeper understanding of these dynamics is crucial for establishing robust monitoring frameworks and implementing targeted strategies to mitigate the public health and environmental risks posed by AMR or potential pathogens in FN communities.

## Materials and Methods

### Sample Site, Collection, and Processing

Wastewater samples were obtained from an FN community with a population of less than 5,000 people, and situated two hours driving distance from Winnipeg, Manitoba. Permission to collect samples was granted by the OLs manager and operators under community anonymity. Samples were collected over a seven-month period. Samples were collected on the following dates, in chronological order: September 09, 2022 (T1); October 21, 2022 (T2); November 16, 2022 (T3); December 09, 2022 (T4); and February 10, 2023 (T5); March 17, 2023 (T6); and April 28, 2023 (T7).

One-liter samples (n=2) were collected in sterile plastic bottles from surface water and OLs. Treated wastewater effluents are released into the Assiniboine River. During each sampling event, a control sample was collected UP from the outfall/discharge site from the OLs and the community. Wastewater samples included those from RS, lagoons (LG), SAGR, and UV-treated sample or effluent (EF). Metadata such as water, dissolved oxygen, conductivity, and atmospheric pressure were measured during sample collection using an Orion Star A323 DO/RDO probe (Thermo Fisher Scientific, Waltham, MA, USA). A Hanna HI98127 pHep4 pH/ temperature tester (Smithfield, RI, USA) was used to assess pH and temperature. Samples were transported to the laboratory using ice packs and coolers and stored at 4°C until further processing.

Within 24 hours of collection, heterotrophic (HC), total coliform (TC), and *Escherichia coli* (EC) plate counts were assessed on the samples using Chromocult coliform agar (Sigma-Aldrich, Oakville, CA, USA) following the manufacturer’s instructions. For UP, SAGR, and EF samples, 100 μL of the homogenized samples were directly spread onto a Chromocult plate and incubated at 37 °C for 24 hours (10). For RS and LG, samples were diluted tenfold, and 100 μL of the diluted sample was plated and incubated as described earlier. Negative controls included the spread of 100 μL of Milli-Q water onto Chromocult plates after each event. Plating was performed in duplicate, and after 24 hours, colony counts were recorded. Dark blue/purple colonies that grew on this media were considered presumptive EC. On the other hand, pink to red colonies, plus dark blue/purple, were considered TC. Colonies that grew white, colorless, creamy, or turquoise were added to TC counts and reported as HC counts.

The strategy for isolating and building these libraries was based on established procedures for typing isolates from FoodNet Canada (11), which identify bacterial isolates found in water, genetically related to human, animal, or food samples. The colonies were picked based on the color they grew on Chromocult. Either dark blue/purple or pink colonies were preferentially selected over other colors. Subsequently, two randomly picked colonies were selected per location in each sampling event, re-streaked onto fresh coliform agar plates, and incubated at 37 °C for 24 hours. This transfer process was repeated three times to ensure the isolation of pure colonies. Following the third transfer, each isolate was transferred to a separate 2-mL sterile cryogenic vial and stored in aliquots of lysogeny broth (LB) medium and 25% (v/v) glycerol at - 80 °C for further characterization.

### Bacterial DNA Extraction

Bacterial isolates from the cryogenic vials as described above, were re-grown overnight at 37 °C at constant agitation in a sterile 15 ml tube, containing 5 mL of LB. Following incubation, tubes were centrifuged for 10 min at 3,000 g and 4 °C. Supernatants were discarded, bacterial pellets were resuspended in 300 μL of Tris-EDTA buffer (pH 8.0) and transferred into sterile 1.5 microcentrifuge tubes for DNA extraction. To these tubes, 1 μL of 5 mg/mL RNase A (LGC Biosearch Technologies, Alexandria, MN, USA) as well as 2 μL of Lysozyme solution (LGC Biosearch Technologies, Alexandria, MN, USA) at ∼30,000 U/μL were added, followed by incubation at 37°C for 30 minutes. After this incubation, the reaction was stopped by adding 2 μL of 0.5 M EDTA (Invitrogen, Waltham, MA, USA), followed by a 10-minute incubation at 65°C.

High-molecular weight DNA was extracted using MasterPure Complete DNA and RNA purification kit (LGC Biosearch Technologies, Alexandria, MN, USA) by following the application notes from the manufacturer on purification of soil metagenomic DNA. DNA quantification was assessed using the Qubit fluorometer 3.0 and Qubit 1x dsDNA HS assay kit (Thermo Fisher Scientific, Waltham, MA, USA) as well as the NanoDrop™ Lite Spectrophotometer (Thermo Fisher Scientific, Wilmington, DE, USA), following the manufacturer’s instructions.

### Library Preparation and Sequencing

Isolates were sequenced using Oxford Nanopore Technologies (ONT). Library preparation was conducted using the ligation-sequencing gDNA protocol and the Native Barcoding Kit 24 V14 from ONT (Oxford, UK). The protocol was followed without any modifications, and sequencing was performed using the MinION sequencer (ONT) with R10.4.1 flow cells (FLO-MIN114) and MinKNOW software (Version 23.11.7). Approximately 400 ng of each DNA sample was utilized for library construction. Before the run, a flow cell check was conducted via the MinKNOW software, which indicated around 1000 pores available for sequencing. Each run contained 12-15 samples multiplexed on each flow cell. A volume of 75 μL from the pooled library was loaded into the flow cell and sequenced for 72 hours. Base calling and demultiplexing of barcoded reads were executed using the MinKNOW software, with all other parameters set to default values. Whole genome sequences from the isolates are available in the NCBI genome datasets (SRA) under Bioproject ID: PRJNA1129067.

### Data analysis

The generated FastQ files from the MinKNOW software were analyzed using the Geneious Prime platform (11.0.18+10) (https://www.geneious.com). Furthermore, genomes for each isolate were assembled using the de-novo assembly option. The assembly method used Flye (12) with input data type set at Nanopore error corrected reads of <2%. Isolates with at least 30X coverage were included in the analysis. For the context of genome identification, 4 online servers were used: TYGS (13), PubMLST (14), RAST (15), and MiGA(16). Additionally, an analysis of ARGs from the assemblies was performed using the Comprehensive Antibiotic Resistance Database (CARD) (17). The specifications employed in the ARGs analysis encompassed examination under the “RGI main” tab, utilizing specific criteria for the DNA sequences. These criteria were delineated by the parameters of perfect, strict, and loose hits. The analytical process excluded the application of the “nudge” parameter and maintained a focus on high-quality coverage. All outcomes derived from the taxonomy tools and CARD analysis were compiled and organized within Excel spreadsheets. In this context, the data were systematically sorted and included to parameters such as consensus strain ID (across taxonomy servers), cut-off values, sample sites, time point, Contig information, AMR family classifications, and associated antibiotic drugs. A bit-score of 50 was used as a cut-off for inferring homology and presenting ARGs (18).

To assess the origin of the ARGs (whether chromosome, plasmid, or phage sequences), we used the SourceFinder tool (19). This integration of information facilitated examination of the genetic and contextual attributes of each sample. HC, TC, and EC counts were analyzed using a Kruskal-Wallis one-way ANOVA. Preliminary statistical tests between factors such as time and site/treatment included Friedman’s two-way nonparametric ANOVA with a randomized complete block design. Statistical analyses were performed using the Statistical Analysis System (SAS, version 9.4 for Windows). A *p*-value of 0.05 was assumed for all tests as a minimum level of significance.

## Results and Discussion

### Bacterial counts

Friedman’s two-way nonparametric ANOVA did not detect significant differences (p>0.0631) for time as variable for any of the HC, TC, or EC counts. Site/treatment was detected as a significant variable (p≤0.0056) for the colony counts. Average numbers of 1.14 x 10^6^, 4.37 x 10^5^, and 6.92 x 10^2^ CFU/ml of sample were observed for HC, TC, and EC, respectively.

As expected, the highest HT and TC counts were detected in RS (p≤0.0023) compared to UP, SAGR or EF (Figure 1). Bacterial counts significantly decreased post SAGR treatment. No other significant differences were detected for the HT and EC colony counts among sites such as SAGR, EF or UP. It is important to note that EC counts were variable across sampling sites and mostly detected in RS, the site with a relatively high-water temperature (11.8 ± 7.5 °C) compared to the other locations. Table 1 summarizes the water quality parameters for the locations analyzed in this study.

**Figure 1.**
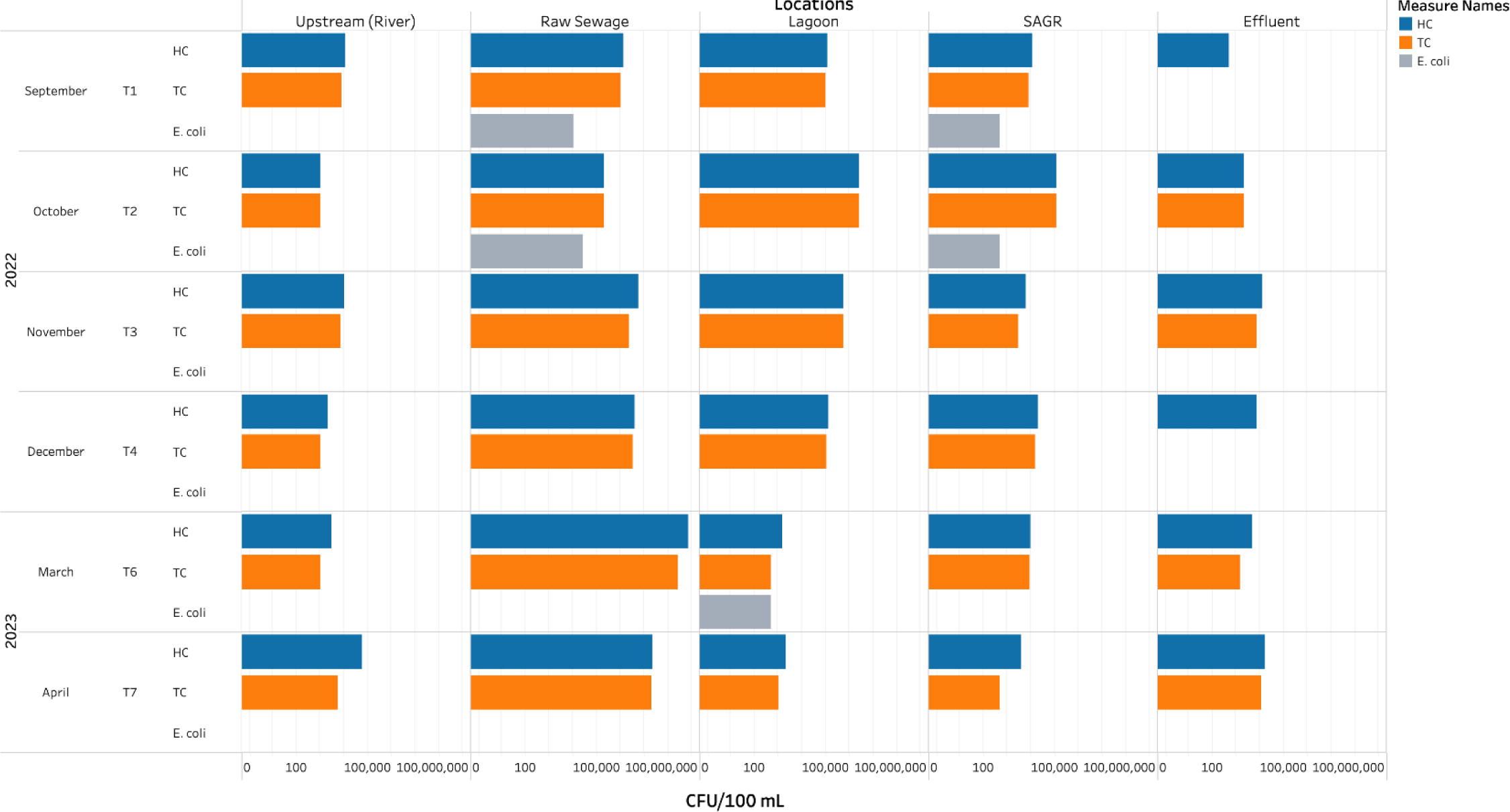
Representation of bacterial counts (Heterotrophic= HC, Total Coliforms = TC, and *Escherichia coli* =E. coli) collected from the various locations and time series assessed in this study. Upstream (river) and oxidation lagoon treatments, including raw sewage, lagoon, Submerged Attached Growth Reactor =SAGR, and effluent (after UV irradiation) are depicted. HC, TC, and EC had on average 1.14 x 10^6^, 4.37 x 10^5^, and 6.92 x 10^2^ CFU/ml of sample, respectively. Bacterial counts differed significantly across sites (p≤0.0056). Raw sewage exhibited the highest HC and TC counts encountered.

**Table 1.** The average water quality parameters assessed from September 2022 to April 2023 in the oxidation lagoons and upstream (river water) near a First Nation community located in Manitoba.

| Sites | DO (mg/L) | Temperature (°C) | pH | Atmospheric pressure (mmHg) |
| --- | --- | --- | --- | --- |
| Upstream<br>(Control) | 6.4 ± 5.4 | 6.2 ± 8.5 | 9.3 ± 0.5 | 733.6 ± 5.9 |
| Raw Sewage | 7.3 ± 4.8 | 11.8 ± 7.5 | 8.1 ± 0.2 | 732.0 ± 8.7 |
| Lagoon | 10.4 ± 2.3 | 9.7 ± 7.3 | 8.2 ± 0.2 | 731.2 ± 6.7 |
| Submerged<br>Attached Growth<br>Reactor | 10.8 ± 2.0 | 9.6 ± 7.4 | 8.3 ± 0.3 | 731.3 ± 6.3 |
| Effluent (UV) | 10.3 ± 1.4 | 9.8 ± 6.7 | 8.2 ± 0.3 | 730.6 ± 6.0 |

### Whole genome sequencing

A total of 58 isolates were found to be distributed among 11 genera. Although 70 isolates were originally stored, not all of them regrew or were viable. At least one isolate was sequenced per sample and time point. The 58 isolates distributed among 11 genera, belonged to 3 families: Moraxellaceae (*Acinetobacter*); Aeromonadaceae (*Aeromonas*); and *Enterobacteriaceae* (*Buttiauxella*, *Citrobacter*, *Enterobacter*, *Escherichia*, *Lelliottia*, *Pantoea*, *Rahnella*, *Serratia*, and *Yersinia*). It is important to mention that recent taxonomic classification has included *Rahnella, Serratia, Yersinia* in the family Yersiniaceae, while *Pantoea* has been reclassified as Erwiniaceae (20, 21). As described in the materials and methods section, Chromocult coliform agar was used following the manufacturer’s instructions for monitoring *Enterobacteriaceae*, specifically TC and *E. coli* (Sigma-Aldrich, Oakville, CA, USA). Despite this, different genera such as *Acinetobacter* (*Moraxellaceae*) and *Aeromonas* (*Aeromonadaceae*) were isolated and further identified. The presence of background microorganisms in chromogenic coliform media poses a challenge that could have been addressed by adding vancomycin and cefsulodin (22). In the present study, we followed the manufacturer’s instructions, which contributed to the isolation and further sequencing of the background colonies. It is worth noting that plates inoculated with Milli-Q water did not yield any colonies.

The most abundant bacterial genera found in all sampling sites was *Aeromonas* (29/58 or 50%), followed by Serratia (9/58 or 15.5%); *Pantoea* and *Escherichia* (4/58 or 7.01% each); *Lelliottia* (3/58 or 5.26%), *Rahnella*, *Enterobacter* and *Buttiauxella* (2/58 or 3.50% each), and finally, *Acinetobacter*, *Yersinia* and *Citrobacter* (1/58 or 1.75% each). Genus *Aeromonas* was prevalent in each location (Figure 2), which is in agreement with previous studies, as members of the *Aeromonas* genus are reported to be ubiquitous in aquatic environments such as freshwater and sewage (23, 24). In this study, 7 out of the 11 genera were observed in RS and included *Aeromonas*, *Citrobacter*, *Enterobacter*, *Escherichia*, *Lelliottia, Rahnella*, and *Serratia*. These findings on RS are not surprising, as sewage harbors a significant richness of microbes. On the other hand, treatments such as LG and SAGR had the least number of genera with only 3 each. Both LG and SAGR contained *Aeromonas* and *Serratia*, but differed in their third genus. *Rahnella* was present in LG, while *Escherichia* was present in SAGR. EF contained 5 out of the 11 genera including *Aeromonas*, *Buttiauxella, Serratia, Acinetobacter,* and *Escherichia.* Finally, control location UP also revealed 5 out of 11 genera: *Aeromonas, Pantoea. Serratia, Enterobacter,* and *Yersinia.* The bacterial counts (Figure 1) and the lower diversity of live isolates (Figure 2) in SAGR and EF suggest a decline in microbial richness. A great number of OLs in rural areas, including FN reserves, only rely on primary physical treatment (3). The OLs here studied employ two additional treatment stages, such as SAGR and UV-treatment for disinfection, which seemed to effectively reduce the bacterial load.

**Figure 2.**
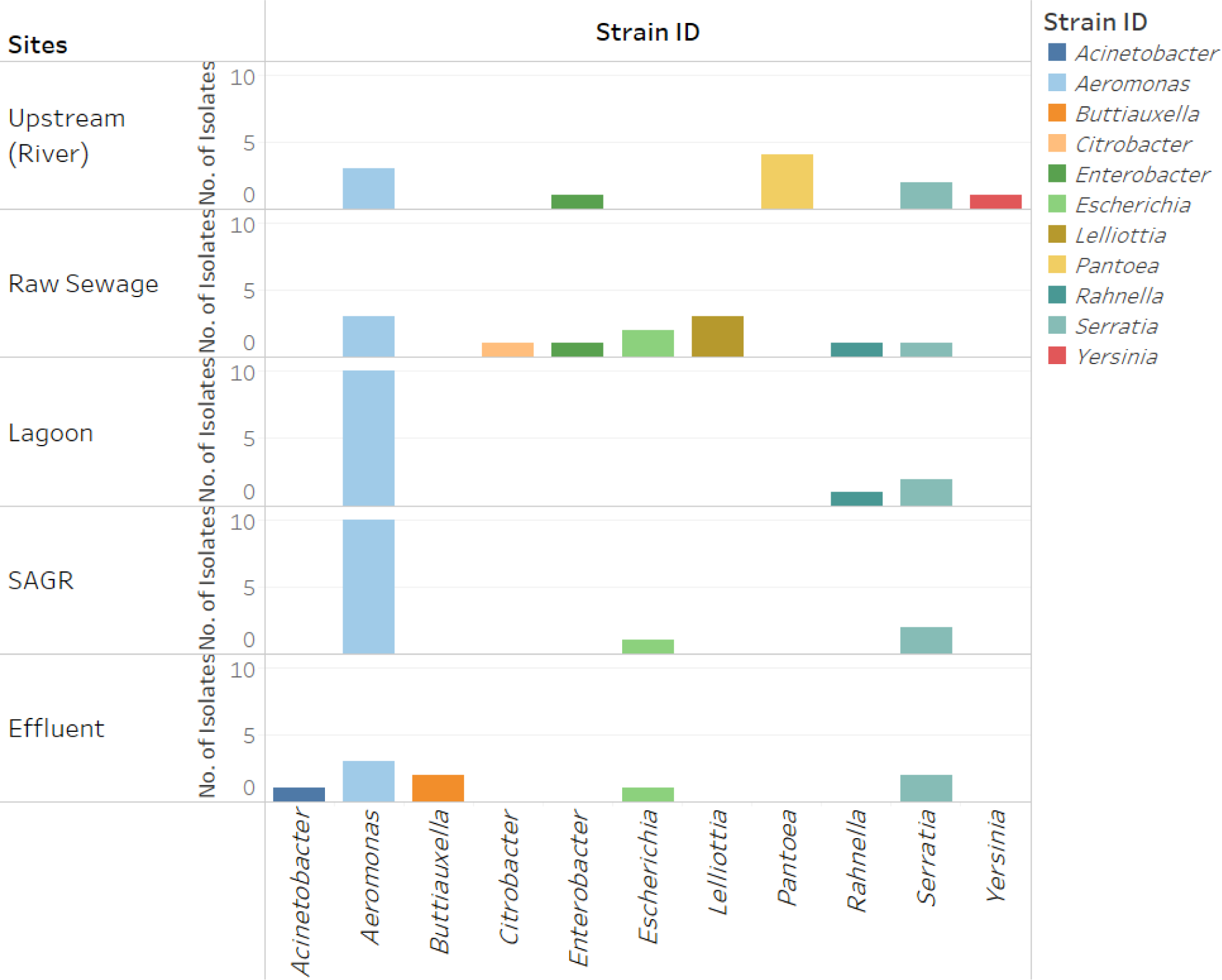
Graphic representation of the number of bacterial isolates and genera found at the various sampling locations of the Oxidation Lagoons (OLs) located in a First Nation community in Manitoba. Submerged Attached Growth Reactor =SAGR. Genus *Aeromonas* was prevalent in each location. Bacterial diversity shows a decrease following SAGR and UV treatment (effluent).

### Antibiotic Resistance Mobilome in Isolates

A total of 32,559 elements distributed in 32134 loose, 397 strict, and 28 perfect hits were classified by CARD as ARGs. Supplemental Table 1 includes the complete information of ARG observed in the chromosome and MGEs, such as plasmids and phages. A total of 302 MGEs (or 0.93% of the total ARGs) were found to be associated with 8 strains including: *Aeromonas*, *Enterobacter*, *Escherichia*, *Lelliottia*, *Pantoea*, *Rahnella*, *Serratia* and *Yersinia*. MGEs were not detected in *Acinetobacter*, *Buttiauxella,* or *Citrobacter* strains. Figure 3 depicts the AMR gene families associated with these MGEs. ARGs detected in plasmids and phages represented 91.4% and 8.60%, respectively. No mobile genetic elements were observed in the isolates for all locations during September 2022.

**Figure 3:**
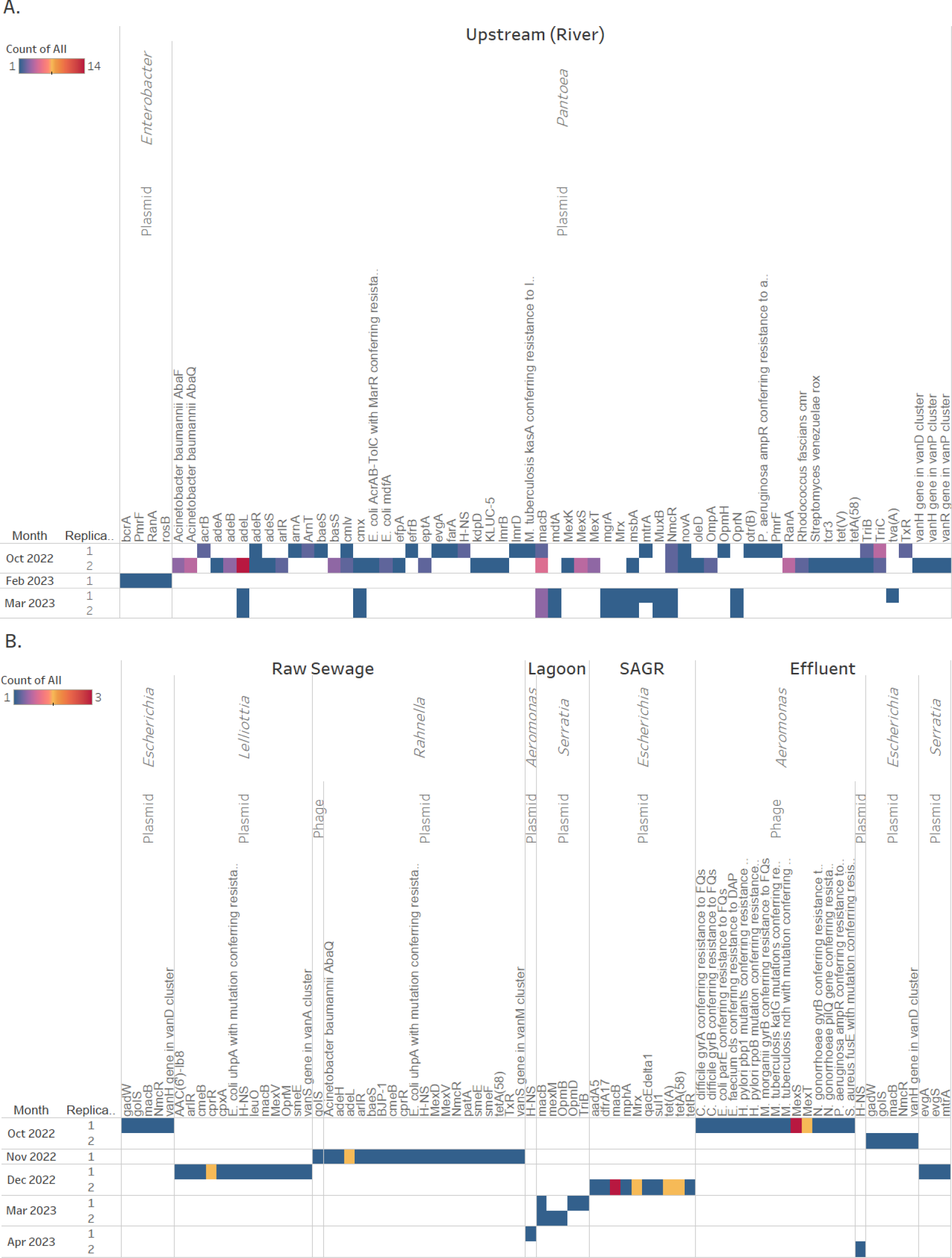
Antimicrobial resistance (AMR) gene families associated with the different mobile genetic elements (MGE) encountered in different bacterial strains identified in the various locations of the Oxidation Lagoons (OLs) located in a First Nation community in Manitoba. Upstream (river sample) was also included in the analysis. A) AMR genes observed in bacterial isolates from the Upstream site. B) AMR genes encountered in bacterial isolates from the OLs treatments, including raw sewage, lagoon, Submerged Attached Growth Reactor =SAGR, and effluent (after UV irradiation). A bit score of 50 was used as a cut-off for inferring homology and presenting AMR genes.

The majority of plasmid-mediated ARGs (187/276, equivalent to 67.8%) were detected at the UP location and distributed mainly in *Pantoea* strains (180/276) compared to *Yersinia* (1/276) and *Enterobacter* (6/276) strains. These ARGs (70/180 ARGs) were associated mainly with efflux pumps genes such as resistance-nodulation-cell division (RND) pumps that confer resistance to multiple antibiotics, including macrolides, β-lactams, tetracyclines, and fluoroquinolones, among others (25, 26). Evidence of plasmid-encoded RND efflux pumps conferring resistance to multiple antibiotics has been reported recently (27, 28)

Other genes that encode for putative efflux pumps detected in *Pantoea* strains were ATP-binding cassette (ABC) antibiotic efflux pumps (33/180 ARGs) and major facilitator superfamily (MFS) antibiotic efflux pump (30/180 ARGs) (Figure 3 and Supplemental Table 1). ARGs detected in the plasmid mobilome of *Yersinia* were associated with RND antibiotic efflux pump (1/187 ARGs). On the other hand, efflux systems such as ATP-binding cassette (ABC) antibiotic efflux pumps (3/187 ARGs), major facilitator superfamily (MFS) (1/187), and pmr phosphoethanolamine transferase (2/187) were observed in *Enterobacter*. It is important to mention that the UP location may be influenced by confined animal feeding operations located nearby. MGEs carrying ARGs in *Pantoea* were observed during October 2022 and March 2023. The mobilome profile among *Pantoea* isolates from October 2022 at the UP site differed from that of isolates from March 2023. This genus showed the most significant amount of hits for macB (n=15) and adeL (n=17) genes. macB encodes for an efflux pump that transports over a dozen-membered lactones outside the cell membrane (29). These membered lactones are macrolide inhibitors that bind to the 23S rRNA subunit and block the mechanism that allows for elongation in protein synthesis for bacteria (30). Macrolides are commonly used to treat infections ranging from simple skin infections to sexually transmitted diseases (31). Plasmid-virulence and high abundance of macB have been reported in similar environments for *Enterobacteriaceae* members as well as other genera such as *Acidovorax*, *Hydrogenophaga*, *Methylotenera*, *Dechloromonas*, and *Nitrospira* (29, 32). These findings highlight plasmid-encoded macB gene crossing families in waters impacted by wastewater effluents. On the other hand, adeL is a regulator for the AdeFGH operon, which encodes for an efflux pump conferring resistance to tetracyclines and fluoroquinolones, and reported in *A. baumannii* plasmids (33). Point mutation in the adeL gene could lead to overexpression of AdeFGH operon and thus would express a phenotype of high-level resistance to clindamycin (26). It has also led to additional mechanisms for high-level resistance to fluoroquinolones and decreased susceptibility to tigecycline, the first drug in the glycylcycline class of antibiotics used to treat certain serious bacterial infections (26, 34). Genes NmcR and TriC also showed significant hit counts with n=10 and n=6 in total, respectively. NmcR, commonly found in *Enterobacter cloacae*, regulates NmcA β-lactamase, which is phenotypically expressed as resistance to carbapenems and β-lactamase inducibility (35, 36). While there are still options for Gram-positive bacteria resistant to carbapenem and β-lactamase, Gram-negative resistant bacteria against these antibiotics are treated with more last-resort drugs such as polymyxins and fosfomycin (37, 38). Studies on TriC in *Pseudomonas aeruginosa* have shown that this gene is part of the TriABC efflux pump system, which expresses a phenotype for triclosan resistance. (39). Triclosan is a very common antiseptic agent used in a variety of cleaning products as well as soaps and body washes (40). In this context, TriC and MexK genes, both contributors of resistance to triclosan, were present in the plasmid of *Pantoea* isolated in October 2022. Often NmcR and TriC genes are found in the chromosomes of non-pathogenic species from aquatic environments, evidence has found them in conjugative plasmids or pathogenic species (35). Although *Pantoea* species rarely cause disease in healthy individuals, they have been reported to impact immunocompromised patients in nosocomial or agricultural settings (41–43). As previously indicated, it is important to point out that the UP location is affected by discharges from nearby confined animal feeding operations for poultry, pork, and beef cattle, which are unrelated to the FN community. Plasmids or other MGEs may have been disseminated to the UP location through horizontal gene transfer and could have influenced the genes identified. The inventory of ARGs detected in RS was broader compared to other processes within the OLs. The most representative plasmid-mediated ARGs included RND (20/54) and MFS (12/54) pumps, which together accounted for 32/54 (59.3%) of the AMR families found. The ARGs encountered in RS were distributed among *Rahnella, Lelliottia, Escherichia*, and *Enterobacter* strains, with *Rahnella* having the highest counts of RND (12/20) and MFS (6/12) pumps, respectively. Moreover, the RS location was one of two places where phage-like structures associated with AMR genes were identified (EF also reported phages).

Analysis of ARGs in phages detected by CARD in RS documented the presence of RND pumps in *Rahnella* during T3 (Supplemental Table 1). *Rahnella spp*. is a gram-negative rod that was isolated from freshwater in the 1970s and has been occasionally associated with human diseases in immunosuppressed individuals (44). A total of five plasmids and one phage-like element carrying ARGs were detected in *Rahnella* isolates. Within this context, *Rahnella* isolates have been reported to harbor multiple plasmids (45, 46). In fact, the chromosomally encoded β-lactamase found in *Rahnella* is suggested to be a progenitor of the plasmid-encoded β-lactamases found in other bacteria. (47, 48). The presence of these MGEs in *Rahnella*, along with recent studies identifying multiple ARGs in these species, supports the evidence of their antibiotic resistance capacity (53). Studies on the proteomics of bacteria-phage interactions have identified several bacterial defense proteins against phages, including multidrug efflux RND transporters, porins, and RM system type I methyltransferase, among others(49). Interestingly, the golS gene, a regulator activated in the presence of golD and promoter of expression of efflux pump, was observed in the phage-like entity in this location. These entities, also known as gene transfer agents (GTAs), have been reported to contain random pieces of the host cell (50). Although golD was not detected in this GTA, this random transfer among prokaryotes is considered influential in horizontal gene transfer (51). While current literature indicates that ARGs are rarely directly encoded in phage genomes (52), these findings suggest that phage-derived ARG transduction should not be dismissed.

Similar to RS, the most common plasmid-mediated efflux pump genes in LG were RND pumps (6/9) distributed among *Serratia* strains. Other plasmid-mediated bacterial efflux pumps identified in smaller proportions in LG included ABC (2/9) pumps in *Serratia* and MFS (1/9) in *Aeromonas.* The *Serratia* genus comprises members recognized as opportunistic nosocomial pathogens capable of causing various infections in humans (53). Previous studies have also found genes associated with efflux pump systems in *Serratia* plasmids (53). The most dominant gene in this location was macB which as previously mentioned, plays a role in extruding antibiotics and export virulence factors from Gram-negative bacteria (29, 54). Research has reported the presence of the MacAB efflux pumps in *Serratia marcescens*, emphasizing the role of these pumps in protecting the bacteria from aminoglycoside antibiotics, polymyxins, and peroxide-mediated killing (55). Identifying the resistome and mobilome of these pumps could aid in finding plausible targets for inhibition, thereby improving the effectiveness of antibiotic therapy.

In SAGR, only MGEs from plasmids were reported. The most representative genes associated with efflux systems were MFS (7/17) and ABC (4/17) pumps dispersed in *Escherichia* strains. These transport systems have been proposed to maintain phenotypic tolerance of drugs, and possibly other environmental stresses, in *Escherichia coli* (56). tetA and tetR genes, responsible for conferring tetracycline resistance (57) as well as the previously mentioned macB, were some of the most relevant genes found in plasmids of *Escherichia* (Supplemental Table 1) Plasmid-mediated tetracycline resistance encoding genes have previously been isolated from multi-drug resistant *E. coli* in environmental samples (58). Despite an overall decrease in bacterial richness at this location, we witnessed the reappearance of *Escherichia* strains (not identified in the LG stage) (Figure 2). Except for lagoon treatment, MGEs carrying ARGs were observed in *Escherichia isolates* (Figure 3B) in all wastewater treatment processes. This fluctuation can be attributed to the fact that samples collected at each stage of the OL were distinct, representing different treatment periods. The retention time (the mean time of wastewater residence in lagoon stages) in this OL was approximately 10 ± 4 days in the lagoons and 9± 1 days in SAGR according to OL operators, which further underscores the variability between sampling points and the dynamic nature of the microbial communities. Consequently, it is not feasible to conclude a consistent survivability pattern of a specific bacteria family based on these cross-sectional samples.

Regarding EF, the records of plasmid-mediated efflux pumps genes were low, accounting for only 1.3% of the total MGEs (13 out of 1,724), when compared to other sites (Figure 3). The most representative plasmid-mediated AMR families were MFS (3/9) pumps reported in *Serratia* and *Aeromonas,* as well as RND (3/9) encountered in *Escherichia* and *Serratia.* Although SourceFinder classified these contigs as plasmids, it is important to note that the presence of prophages in whole genome sequences may contribute to the misclassification of these elements (23). For instance, additional analysis of contigs using RAST and Subsystems (results not shown) detected capsid, scaffold, and terminase phage and mobile element proteins. Furthermore, in this location, a more complete set of ARGs present in phage sequences (47.2 Kb) was detected in *Aeromonas* strains (Figure 3 and Supplemental Table 1). The most commonly identified phage-mediated efflux pump genes were RND (12/26 or 46.15%), which confer resistance to fluoroquinolones, diaminopyrimidine, and phenicol antibiotics. Other phage-mediated genes found in smaller proportions in *Aeromonas* were fluoroquinolone resistant gyrA, parE and fluoroquinolone and zoliflodacin resistant gyrB (Figure 3 and Supplemental Table 1). In this context, evidence of horizontal gene transfer in quinolones resistance-determining regions of gyrA, gyrB, and parE have been reported in Gram-negative bacteria (59, 60). Further annotation using RAST, blastn, and PHASTER identified these sequences as Enterobacteria phages, *Aeromonas* phage vB_AsaM-56, and others classified within the class Caudoviricetes as well. As previously reported, genes conferring reduced susceptibility to fluoroquinolones have been found in enterobacteria phage fractions, specifically in somatic coliphages (61). There are only a limited number of sequenced phage genomes reported in the literature, and evidence of these bacteriophages carrying genes that encode putative ARGs is relatively scarce (62). Despite this, these results contribute to the growing evidence supporting the role of phages as potential vehicles for horizontal gene transfer.

### Bacterial diversity

Overall, there was a significant reduction in bacterial counts and isolates from RS to the final UV-treated effluent (Figures 1 and 2). The varying retention times across the OL stages (10 ± 4 days) and the subsequent SAGR system (9 ± 1 day) in addition to more extended retention periods due to winter (frozen OL, sedimentation was slowed) (63, OL main operator, personal communication, 2023) demonstrate that samples from each stage do not represent the same initial wastewater input. This leads to potential variations in microbial composition over time. The variability in *E. coli* counts across treatment stages was notable. At certain time points (e.g., September and October 2022), the OL exhibited minimal or undetectable *E. coli* levels, while downstream stages such as SAGR showed significantly higher counts. This disparity may be attributed to differences in retention times within the wastewater treatment process. Lagoons prior to SAGR can retain wastewater for up to 90 days, making the total residence time from RS input to effluent discharge range from 23 to over 90 days (63, OL main operator, personal communication, 2023). Such variation in retention, coupled with fluctuations in the incoming sewage load and transportation dynamics, can significantly influence microbial diversity and *E. coli* abundance across the various stages. Consequently, the interpretation and comparison of microbiological data across different treatment stages should be conducted cautiously, as samples collected at different stages represent distinct batches of treated wastewater. OLs are prevalent in small communities in Canada, and these systems are relatively affordable to build and operate (2, 64). Lagoon-based treatments offer valuable aid in protecting stream quality by generating effluents with minimal environmental impact. Despite this, studies have reported that the removal efficiency of ARGs by conventional biological water treatment processes is low (65). WWTPs and lagoon-treatment facilities promote optimal conditions for bacterial growth and proliferation, including the provision of nutrients, suitable temperature, and a suitable pH level (66).

Even when treatments with other disinfectants reduce bacterial viability, they do not degrade ARGs. Microbial cells that survive the treatment have been reported to uptake ARGs and other genes from recently killed bacteria (65, 67). Additionally, the selective pressure exerted by antibiotic residues facilitates the proliferation of ARB and the formation of multidrug resistance (66, 68). As displayed in Figures 2 and 3 there is evidence of bacteria that have flown through the UV with ARGs, which follows the growing evidence that sustains that treated effluents transport ARBs and ARGs to the downstream water environment (68). Despite the known capacity of UV radiation to damage bacterial DNA and prevent replication(69), our findings demonstrate a restricted impact of UV treatment on the elimination of ARGs and ARB in the effluent. This limitation is likely due to the UV dose applied in the OLs being insufficient to effectively inactivate all ARB or degrade ARGs, given that the higher doses necessary for such outcomes are generally considered impractical for sustainable wastewater treatment (66). In this context, OLs may represent an important route for AMR dissemination into the environment.

## Conclusions

This work represents an example of isolates observed in OLs, the most common treatment method used by rural communities in Canada, including FN. The facility here studied uses two additional wastewater treatment processes not included in conventional lagoon treatment, such as SAGR and UV. While the OLs treatment reduced overall bacterial loads, this analysis revealed the prevalence of bacteria, resistome, and mobilome distributed throughout the sampling sites (2). MGEs carrying ARGs were detected across all sampled locations. Interestingly, UP exhibited a higher richness of ARGs compared to the EF location. This finding evidences the effectiveness of this OLs treatment in ameliorating contamination impact in the environment, due to the lower ARG richness observed in the EF compared to the reference control site. Plasmids were found to harbor genes that encoded for different efflux pumps, including RND family transporters, the most frequently occurring pumps throughout the isolates. A notable prevalence of the plasmid-mediated genes macB, adeI, tetA, and tetRwere observed across a range of bacterial isolates. Importantly, phage-like structures detected in both the RS and EF samples were found to harbor ARGs, such as gols, the fluoroquinolone-resistance genes gyrA and parE, and the gene gyrB, which confer resistance to both fluoroquinolones and zoliflodacin. This evidence strengthens the hypothesis that phages may serve as mobile genetic elements for ARGs, thereby introducing an additional layer of complexity to the mechanisms of AMR dissemination. The resistome and mobilome persisting in the treated effluent underscore the role of OLs as a conduit for the dissemination of AMR into the environment. Future investigations should prioritize the surveillance of the factors contributing to the emergence of ARBs in these rural systems and their implications for community members and the surrounding environment. It is imperative to establish standardized measures, such as establishing regulatory limits for ARGs discharge, and to execute risk assessment plans to identify and quantify ecological and public health risks posed by the dissemination of ARGs in OLs and WWTPs. Finally, it is key to identify and implement treatment technologies for process optimization and ARGs removal in OLs.

## List of abbreviations

AMR: Antimicrobial Resistance
ARB: Antibiotic-Resistant Bacteria
ARGs: Antibiotic Resistance Genes
EC: *Escherichia coli*
EF: Effluent
FN: First Nation
HC: Heterotrophic Counts
MGEs: Mobile Genetic Elements
OL(s): Oxidation Lagoons
RS: Raw Sewage
SAGR: Submerged Attached Growth Reactor
TC: Total Coliforms
UP: Upstream
WWTPs: Wastewater Treatment Plants

## Acknowledgments

This work was funded by the NSERC-DG program (RGPIN-2022-04508) with additional support from the Research Manitoba New Investigator Operating Grant (No. 5385), both awarded to MUD. Indigenous CREATE granted fellowships to RP, JZ, and MM. RP, JZ and AA were supported by Research Manitoba. MM and KZ were funded by the NSERC-DG program. The authors recognize Kumar lab (University of Manitoba) for sequencing guidance and support. Special thanks to operators, and manager from the oxidation lagoon facility as well as FN community members for their help navigating the terrain. This research was conducted at the University of Manitoba following the First Nations principles of Ownership, Control, Access, and Possession and adhering to the Tri-Council policy involving the First Nations, Inuit and Métis communities. The authors acknowledge that the University of Manitoba campuses are located on original lands of Anishinaabeg, Ininewuk, Anisininewuk, Dakota Oyate, and Denesuline, and on the National Homeland of the Red River Métis.

